# A Novel Secretome Rewrites the Immune Response in Viral Acute Respiratory Distress Syndrome

**DOI:** 10.64898/2026.02.06.704035

**Authors:** Daniel Lapuente, Noelia Mendoza Calvo, Carl P. Lehmann, Belén García, Juan Antonio Quintana, Almudena De Gregorio, Kevin Carpio, Javier Galán-Martínez, Angela R. García-Rendueles, Celia Camacho-Toledano, Fernando Oroz-Gonjar, Paula Desportes, Jara Sanz, Pablo Fernández, Juan Carlos De Gregorio, Francesca Sarno, Juan Pedro Lapuente

## Abstract

**Background:** Hyperinflammatory syndromes such as viral acute respiratory distress syndrome (ARDS) demand immunotherapies that are safe and effective at suppressing cytokine storm. As a novel treatment for ARDS, PRS CK STORM is proposed as a next-generation cell-free secretome derived from co-cultures of M2 macrophages and mesenchymal stromal cells.

**Methods:** We performed a double-blind, randomized controlled trial to test the effects of PRS CK STORM in K18-hACE2 transgenic mice infected intranasally with 10^5^ PFU of SARS-CoV-2 BE.1.1. This was complemented by a mechanistic analysis of this secretome using transcriptomic and COX2 enzymatic activity studies, as well as a compositional analysis of its proteomic and miRNA profile.

**Results:** In a lethal SARS-CoV-2 ARDS mouse model, PRS CK STORM significantly improved lung histopathology to a degree on par with corticosteroids, while stimulating angiogenesis in damaged lung tissue. Early and late cytokine profiling showed marked reductions in IFN-γ. In response to PRS CK STORM treatment, transcriptomic analyses in inflamed macrophages revealed robust downregulation of key proinflammatory drivers (MyD88, TRAF6, IKK2, NF-κB, COX2). *In vitro* enzymatic assays confirmed potent, dose-dependent inhibition of COX2, with high inter-batch reproducibility. A compositional analysis revealed this secretome to be rich in anti-inflammatory miRNAs, immune-modulating proteins, and regenerative factors.

**Conclusions:** PRS CK STORM operates as a multi-target immune recalibrator, enabling broad downregulation of pathological inflammation while promoting tissue repair. Its off-the-shelf, GMP-manufactured format ensures reproducibility and scalability, offering a novel resolution pharmacology approach for cytokine storm syndromes originated from ARDS and beyond.

## Introduction

Acute respiratory distress syndrome (ARDS) is a critical form of non-cardiogenic respiratory failure caused by diffuse pulmonary inflammation. Viral pneumonias— particularly SARS-CoV-2 and influenza—are major triggers via dysregulated immune responses, often culminating in a cytokine storm marked by elevated pro-inflammatory cytokines such as interferon gamma (IFN-γ), interleukin-1 beta (IL-1β), and tumor necrosis factor-alpha (TNF-α) (1–3). This hyperinflammatory state not only drives respiratory failure but also systemic organ injury. Even before the COVID-19 pandemic, ARDS represented a global health burden, with incidence rates ranging from ∼10 to ∼79 per 100,000 persons per year. During the pandemic, nearly one-third of hospitalized COVID-19 patients developed ARDS, with mortality in critical cases reaching 40–60%. Survivors frequently suffer long-term pulmonary sequelae, and the syndrome incurs substantial ICU-related costs, underscoring the urgent need for scalable immunotherapies (4,5).

Cytokine storm reflects a breakdown in immune homeostasis. Pathogen-associated molecular patterns (PAMPs) and damage-associated molecular patterns (DAMPs) activate pattern recognition receptors (PRRs), triggering signaling of nuclear factor kappa-light-chain-enhancer of activated B cells (NF-κB) and inflammasome assembly in macrophages and dendritic cells. This results in massive cytokine production and pyroptosis. Interleukin (IL)-6, via signal transducer and activator of transcription 3 (STAT3) and glycoprotein 130 (gp130) signaling, promotes chemokine-driven leukocyte recruitment, while macrophages remain locked in a pro-inflammatory M1-like phenotype, producing TNF-α, IL-1β, and IL-12. Failure of macrophages to transition to the reparative M2 state leads to sustained tissue damage, endothelial dysfunction, and immune dysregulation (6). This cycle is difficult to break, as redundancies within the immune system as well as patient heterogeneity can render single-cytokine blockade treatments insufficient, suggesting the need for combination or multi-target therapies (7). In the context of COVID-19 related ARDS, the standard intervention is treatment with the corticoid dexamethasone, which while modestly effective in reducing patient mortality, is associated with a variety of adverse effects including immunosuppression (8–10).

Physiologically, resolution of inflammation is mediated by feedback circuits involving a variety of components including IL-10, interleukin-1 receptor antagonist (IL-1Ra), soluble TNF receptors, efferocytosis, and pro-resolving mediators. These mechanisms restore immune balance without compromising host defense (11,12). Cell-derived secretomes comprising cytokines, chemokines, extracellular vesicles, and microRNAs (miRNA), have emerged as promising tools to replicate this endogenous multi-faceted resolution program (13–16).

Building on this, we developed PRS CK STORM, a next-generation secretome generated by co-culturing M2-polarized macrophages with mesenchymal stromal cells (MSCs). This model mimics the crosstalk of resolving tissues, enriching the secretome with IL-1Ra, IL-10, hepatocyte growth factor (HGF), and insulin-like growth factor (IGF), while depleting C-C motif chemokine ligand 2 (CCL2) and IL-8. *In vitro*, PRS CK STORM suppresses inflammatory cytokine release comparably to high-dose corticosteroids and is not cytotoxic (17,18). In this present study, we sought to assess its efficacy *in vivo* in the context of ARDS, as well characterize its mechanism of action and composition. We propose PRS CK STORM as a novel biologic with a reproducible, good manufacturing practices (GMP) format that can simultaneously downregulate cytokine storm mediators while promoting lung repair.

## Methods

### *In vivo* SARS-CoV-2 ARDS Model and Treatment

K18-hACE2 transgenic mice were intranasally challenged with 1×10^5^ plaque-forming units (PFU) of SARS-CoV-2 Omicron BA.5 (LD_100). This dose yields a fulminant cytokine storm and near-uniform lethality by day 8 (19). Infected mice were treated intravenously with either PRS CK STORM (40 µL) or dexamethasone (5 mg/kg), starting at 24 hours post-infection. All animals developed systemic illness; no group survived beyond day 8. Clinical deterioration was assessed by measuring body weight and observational monitoring for signs of stress. Tissue was harvested at terminal illness for cytokine assays and lung histopathology. For more details, see online supplementary information.

### Production of PRS CK STORM

The production process for PRS CK STORM is described in detail in (17).

### Quantitative RT-PCR Analysis in Macrophages

M1-polarized Tohoku hospital pediatrics-1 (THP-1) cells were generated as described in (20). These were then stimulated with 10 ng/mL Lipopolysaccharide (LPS) (Sigma– Aldrich) and simultaneously treated with a 1:1 volume ratio of PRS CK STORM or cell culture media (control). Total RNA was extracted and reverse-transcribed using the ThermoFisher cDNA synthesis kit (Ref. 4368814) according to manufacturer’s instructions. Quantitative PCR (qPCR) was performed using validated primers targeting key proinflammatory and immunoregulatory genes. GAPDH was used as the housekeeping gene for normalization, and relative gene expression was calculated using the ΔΔCt method. Primer sequences used for all examined genes are detailed in table S1.

### COX2 Inhibition Analysis

The ability of PRS CK STORM to inhibit cyclooxygenase-2 (COX2) at the enzymatic level was evaluated using a COX2 Inhibitor Screening Kit (Abcam #ab283401) according to manufacturer’s instructions. The assay was challenged with 2-10 µL of PRS CK STORM or Celecoxib, a specific COX2 inhibitor control.

### Statistical Analysis

Statistical analysis for blood capillary quantification, bodyweight, qPCR, and COX2 inhibition assays were performed using one-way ANOVA, with Prism 9 (GraphPad Software Inc.). *In vivo* total lung pathology scores were analyzed using a Tukey HSD post hoc test. P values <0.05 were considered to indicate statistical significance. Statistical analysis of the functional enrichment analysis is described in the online supplementary information.

## Results

### Treatment with PRS CK STORM Preserves Lung Integrity and Facilitates Regeneration in an *In Vivo* SARS-CoV-2 ARDS Model

To validate PRS CK STORM in a severe viral ARDS context, K18-hACE2 transgenic mice were intranasally challenged with 1×10^5^ PFU of SARS-CoV-2 Omicron BA.5 (LD_100), followed by treatment with PRS CK STORM or dexamethasone (corticoid). The aforementioned viral dose yields a fulminant cytokine storm associated with extremely high lethality by day 8 (19). To confirm effective induction of the model, viral load was assessed by analyzing the cycle threshold (Ct) of RT-qPCR from lung homogenates collected at terminal illness. The resulting low Ct values in the positive control and treatment groups, relative to the negative control, confirmed a high viral burden consistent with a fulminant infection (Figure 1A). This is comparable to very severe human COVID-19, where mortality risk reaches ∼35% and 29.1% of survivors require intubation (21). No notable differences in viral load were observed among the three infected groups, indicating PRS CK STORM did not affect viral clearance.

**Figure 1.**
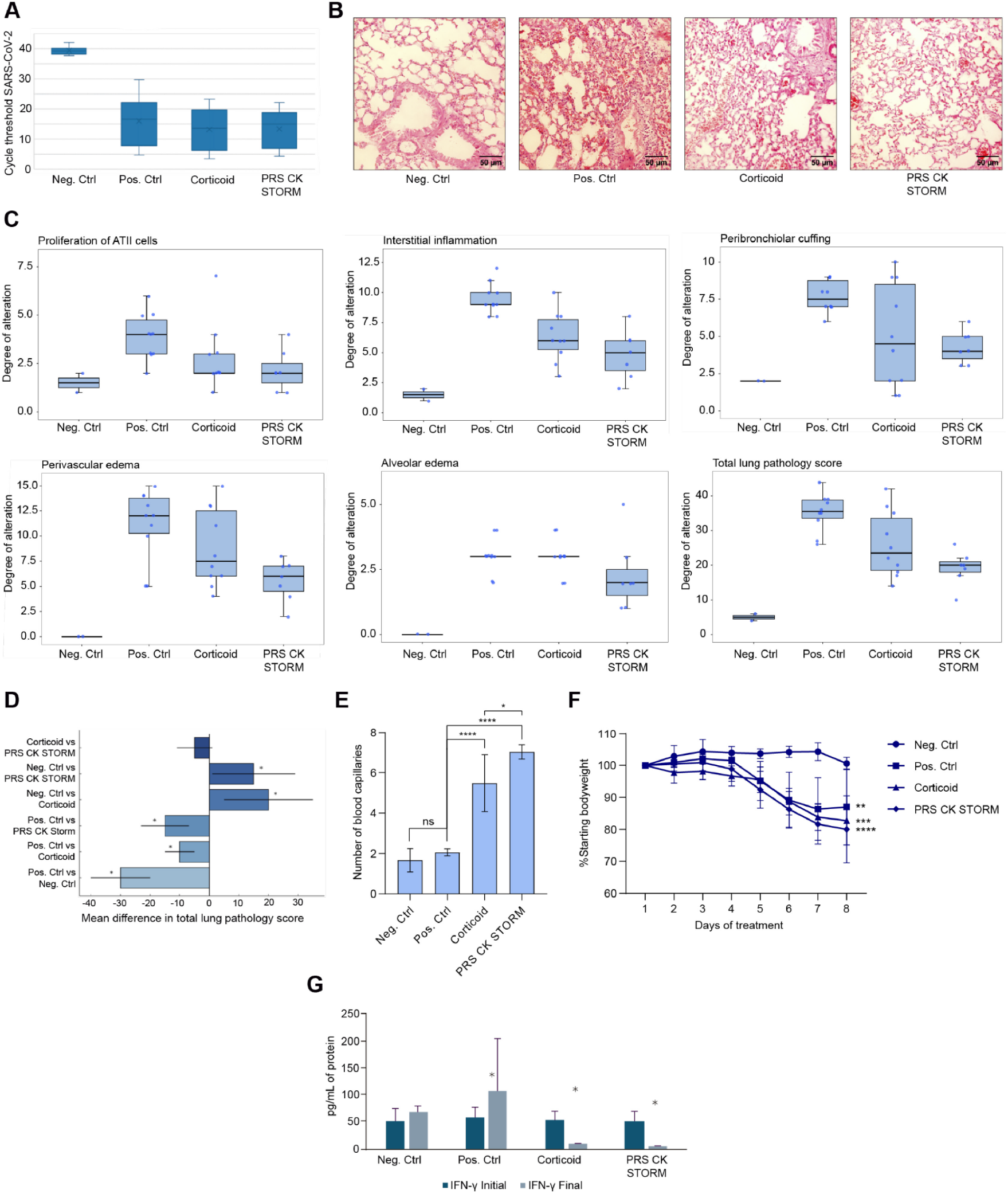
A SARS-CoV-2 ARDS mouse model displays improved lung pathology after treatment with PRS CK STORM. Neg. Ctrl corresponds to uninfected animals; Pos. Ctrl denotes infected, untreated animals; the Corticoid group received corticosteroid (dexamethasone) treatment; and the PRS CK STORM group received the secretome therapy. (**A**) Quantification of SARS-CoV-2 viral RNA by RT-qPCR in lung tissue across treatment groups. Box plots represent Ct (cycle threshold) values from lung homogenates collected post-infection at terminal illness. Higher Ct values indicate lower viral load. Infected animals (Pos. Ctrl, Corticoid, PRS CK STORM) showed significantly reduced Ct values compared to uninfected control (Neg. Ctrl), consistent with robust viral replication. Boxes represent interquartile range (IQR), whiskers show minimum and maximum values, and horizontal lines indicate medians; means are denoted by “×”. (**B**) Representative pulmonary histopathology in the different treatment groups. Brightfield microscopy of hematoxylin-eosin-stained lung sections from experimental mice (20x magnifications). (**C**) Scores for alterations in pulmonary parameters. Four random regions of lung (two per lobe), per mouse, were scored based on the following parameters: proliferation of ATII cells, interstitial inflammation, peribronquiolar cuffing, perivascular edema, alveolar edema on a scale of 0 (no alteration) to 4 (high alteration) (see table S2 for further details). The bottom right graph displays the composite total lung pathology score, combining all aforementioned parameters. (**D**) Post hoc Tukey HSD test displaying pairwise group comparisons of total lung pathology scores. Bars represent mean differences (± 95% confidence interval). (**E**) Quantification of pulmonary blood capillaries. The average of five random regions of lung were quantified per mouse. Shown are the mean ± SD for each group. (**F**) Bodyweight of mice during the study. Mean ± SD of percentage of bodyweight relative to day 1. (**G**) Quantification of serum concentration of murine IFN-γ by multiplex ELISA. Bar graphs depict plasma concentrations (mean + SD) of pro-inflammatory cytokine IFN-γ in each treatment group, measured 1 day after infection shortly after initial treatment (Initial, day 1) and at the experimental endpoint (Final, terminal illness). In all graphs, asterisks (*) denote statistically significant differences: *(p < 0.05), **(p < 0.01), ***(p < 0.001), ****(p < 0.0001).

A blinded histopathological analysis of the lungs was conducted postmortem on all animals using hematoxylin–eosin staining. Four random regions of lung (two per lobe) were scored for each mouse according to a predefined protocol (Table S2). The positive control group exhibited widespread alveolar damage, alveolar edema, and interstitial inflammation, a disease phenotype consistent with ARDS (22,23). Corticosteroid-treated lungs showed moderate improvement with residual inflammation, while PRS CK STORM– treated lungs exhibited preserved alveolar architecture with minimal infiltrates and edema, consistent with reduced overall pathological severity (Figures 1B, C, D). Of note, mice treated with PRS CK STORM, and to a lesser extent corticoids, exhibited a significant increase in the number of pulmonary blood capillaries compared to the untreated positive control (Figure 1E). This is indicative of angiogenesis, which facilitates lung recovery and regeneration in the context of ARDS (23,24).

Mice from all SARS-CoV-2 infected groups exhibited significant weight loss (Figure 1F) and severe clinical symptoms such as hunching, lethargy, and piloerection by the end of the trial, with some needing to be sacrificed on day 7 for humane endpoint. However, it should be noted that the experimenters observed on day 6 (one day after the final day of treatment) that the PRS CK STORM treated mice appeared to be more active than those of the positive control or corticoid groups.

Early and terminal cytokine profiling, measuring plasma concentrations of murine IFN-γ on day 1 (1 day post infection and 1 minute post-treatment) and at terminal illness endpoint, revealed the immunomodulatory effects of PRS CK STORM (Figure 1G). As expected, mice from the uninfected negative control group maintained similar levels of IFN-γ between the start and finish of the study, while mice from the positive control group displayed a significant increase in IFN-γ, consistent with a pro-inflammatory immune response (3). In contrast, mice from both the corticosteroid and PRS CK STORM groups exhibited a significant decrease in IFN-γ levels at the end of the experiment. Altogether, although PRS CK STORM did not improve survival in this lethal model, these data support its capacity to suppress systemic inflammation and reduce pathological severity while favoring lung recovery in the context of ARDS.

### Inflammatory Gene Expression in THP-1 Derived Macrophages is Reduced by PRS CK STORM

To further investigate how PRS CK STORM affects the immune response, we analyzed its effect on macrophages in the context of LPS-induced inflammation. Its immunomodulatory effects on M1-polarized THP-1 cells were studied via *in vitro* transcriptomic analyses of genes involved in inflammatory pathways. We began by studying expression levels of the enzyme COX2, a major contributor to inflammation via its production of prostanoids (25). Treatment of LPS-stimulated THP-1 macrophages with PRS CK STORM showed a time-dependent decrease in *COX2* expression (Figure 2A). While shorter treatment times of 6 or 24 hours had no effect, 48 hours of treatment with PRS CK STORM depleted virtually all *COX2* transcripts, thus suggesting that our research product is a powerful inhibitor of this pathway. We then looked at the effect of PRS CK STORM on other key inflammatory genes that are upstream of COX2, over the course of a 48-hour period (Figure 2B). Downregulation was observed for genes TNF receptor associated factor 6 (*TRAF6*), IκB kinase 2 (*IKK2)*, myeloid differentiation primary response 88 (*MYD88*), NF-κB subunit 1 (*NFKB1*) and again *COX2*, which are all key effectors in the toll-like receptor (TLR)/NF-κB signaling cascade (26). These data support a multi-target immunoregulatory activity for PRS CK STORM.

**Figure 2.**
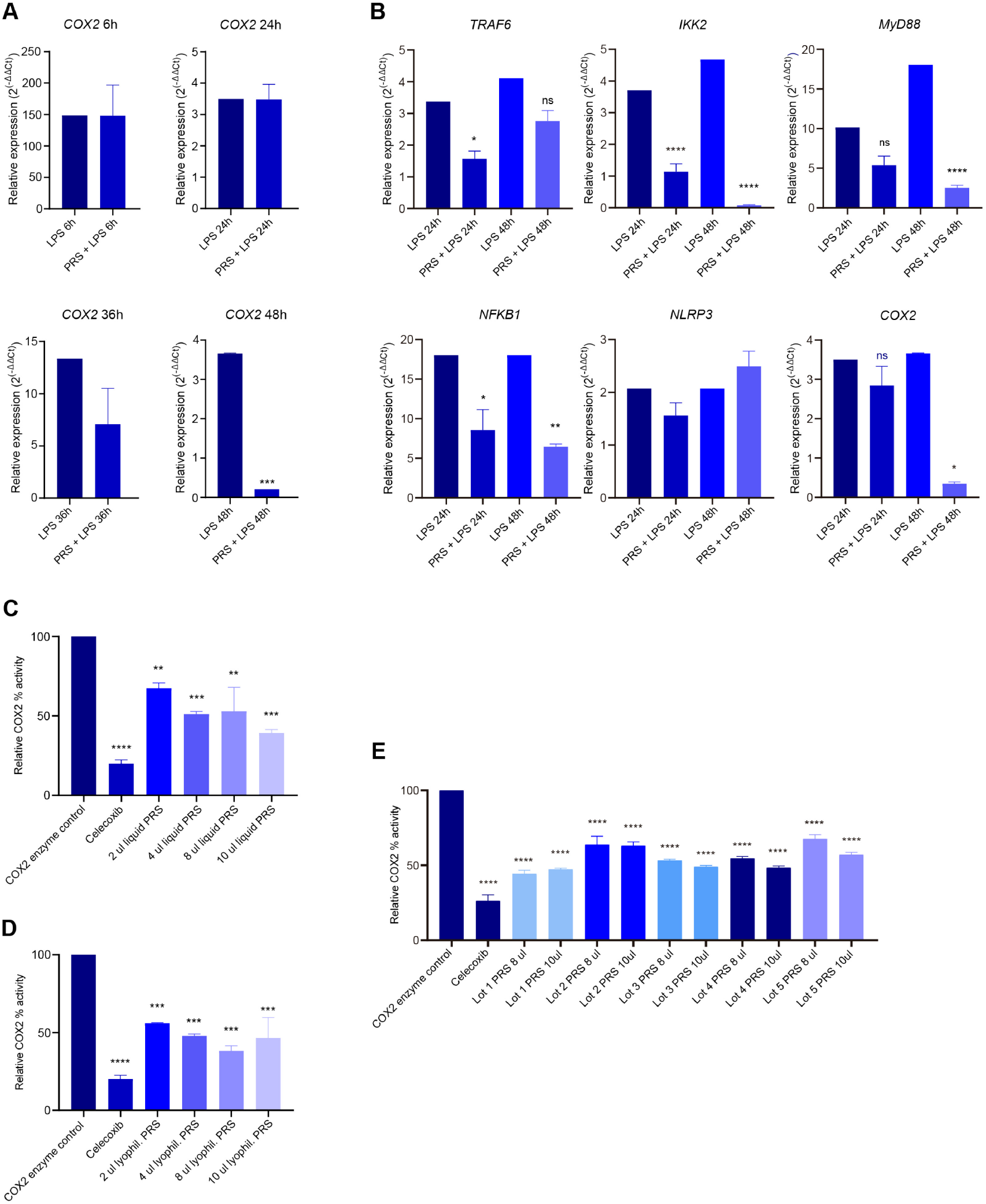
PRS CK STORM inhibits inflammatory gene expression in macrophages and consistently inhibits COX2 enzymatic activity. “PRS” = PRS CK STORM. (**A**) Relative expression of COX2 after 6-, 24-, 36- and 48-hours treatment with PRS CK STORM and LPS. THP-1 derived macrophages were stimulated with 10 ng/ml LPS and simultaneously treated with or without a 1:1 volume ratio of PRS CK STORM for the duration of the experiment. Bar plots display the relative mRNA expression (mean ± SD). Expression was normalized to GAPDH and calculated using the ΔΔCt method. (**B**) Gene expression analysis of pro-inflammatory mediators in LPS-stimulated THP-1 macrophages treated with PRS CK STORM, as in A. Bar plots display the relative mRNA expression (mean ± SD) of TRAF6, IKK2, MYD88, NFKB1, NLRP3, and COX2 at 24 hours and 48 hours post treatment. Expression was normalized to GAPDH and calculated using the ΔΔCt method. (**C**) COX2 enzymatic inhibition by PRS CK STORM in liquid format. COX2 activity was assessed using a fluorometric kinetic assay based on prostaglandin E2 production. The assay was carried out in the absence of inhibitor (COX2 enzyme control), in the presence of celecoxib (inhibition positive control), or with various concentrations of PRS CK STORM in liquid format (2, 4, 8, or 10 µl). The bar plot displays the relative COX2 activity (mean ± SD). (**D**) COX2 enzymatic inhibition by PRS CK STORM in lyophilized format. The assay was as in C, utilizing a lyophilized format of PRS CK STORM rather than a liquid format. (**E**) Inhibition of COX2 activity by PRS CK STORM across five independent GMP batches. The assay was as in C. 8 or 10 µl of each lot of PRS CK STORM was used in the reaction. The bar plot displays the relative COX2 activity (mean ± SD). In all graphs, asterisks (*) denote statistically significant differences: *(p < 0.05), **(p < 0.01), ***(p < 0.001), ****(p < 0.0001).

### PRS CK STORM Inhibits COX2 Activity Directly and Consistently Across Different Production Lots

To determine whether PRS CK STORM acts directly on the COX2 enzyme, an *in vitro* enzyme activity assay was conducted in the presence of PRS CK STORM or celecoxib, a specific COX2 inhibitor that serves as a control. COX2 enzymatic activity was strongly diminished in response to incubation with PRS CK STORM in as dose-dependent manner (Figure 2C), thus demonstrating that this secretome inhibits the COX2 pathway directly. Given that PRS CK STORM is a biologic drug, the possibility of variability between formats and batches was a concern. We repeated the above-mentioned COX2 activity assay to analyze the potency of PRS CK STORM in a lyophilized format (Figure 2D). Here we observed a similar efficacy as compared to that of a liquid format (Figure 2C). We then measured inhibition of COX2 across 5 independent GMP batches of PRS CK STORM, and observed consistent efficacy, with a coefficient of variation of <10% between lots (Figure 2E). These results demonstrate that PRS CK STORM reliably inhibits COX2 enzymatic activity across formats and production lots, supporting its use as a reliable biologic for reducing inflammation.

### Exosomal miRNA Profiling of PRS CK STORM Reveals Dominant Immunoregulatory Signatures

To assess whether its molecular composition reflects its observed therapeutic activity and mechanism of action, we analyzed the presence of potential immunomodulatory elements in PRS CK STORM. As a secretome-based therapeutic derived from co-cultures of M2 macrophages and MSCs, PRS CK STORM is likely enriched in extracellular vesicles, such as exosomes (13,27). We therefore performed exosomal miRNA sequencing. The top 10 most abundant exosomal miRNAs in PRS CK STORM accounted for nearly 80% of the total miRNA content and have all been previously reported to exert beneficial effects in preclinical models of respiratory and inflammatory disease by modulating key pathways involved in immune regulation, inflammation resolution, and tissue repair (Figure 3A and Table S3). A striking feature of this profile was the marked overrepresentation of the let-7 family. Together with hsa-miR-26a; let-7a, b, c, and f have been shown to be significantly downregulated in lung tissue from murine models of ARDS, and inversely associated with expression of pro-apoptotic and oxidative stress-related genes such as superoxide dismutase 2 (*SOD2*) and adenomatosis polyposis coli (*APC)* (28), implicating them in the regulation of injury-associated transcriptional programs. Notably, let-7 has been reported to inhibit the expression of inflammatory cytokines including IL-1β, IL-6, TNF-α, and granulocyte-macrophage colony-stimulating factor (GM-CSF) (29), and negatively regulate PI3K/Akt/COX2 signaling in bacteria-induced pulmonary fibrosis (BIPF) animal models (30). Collectivity, the observed exosomal miRNA profile is consistent with the ability of PRS CK STORM to reduce cytokine levels and lung injury in our *in vivo* ARDS murine model.

**Figure 3.**
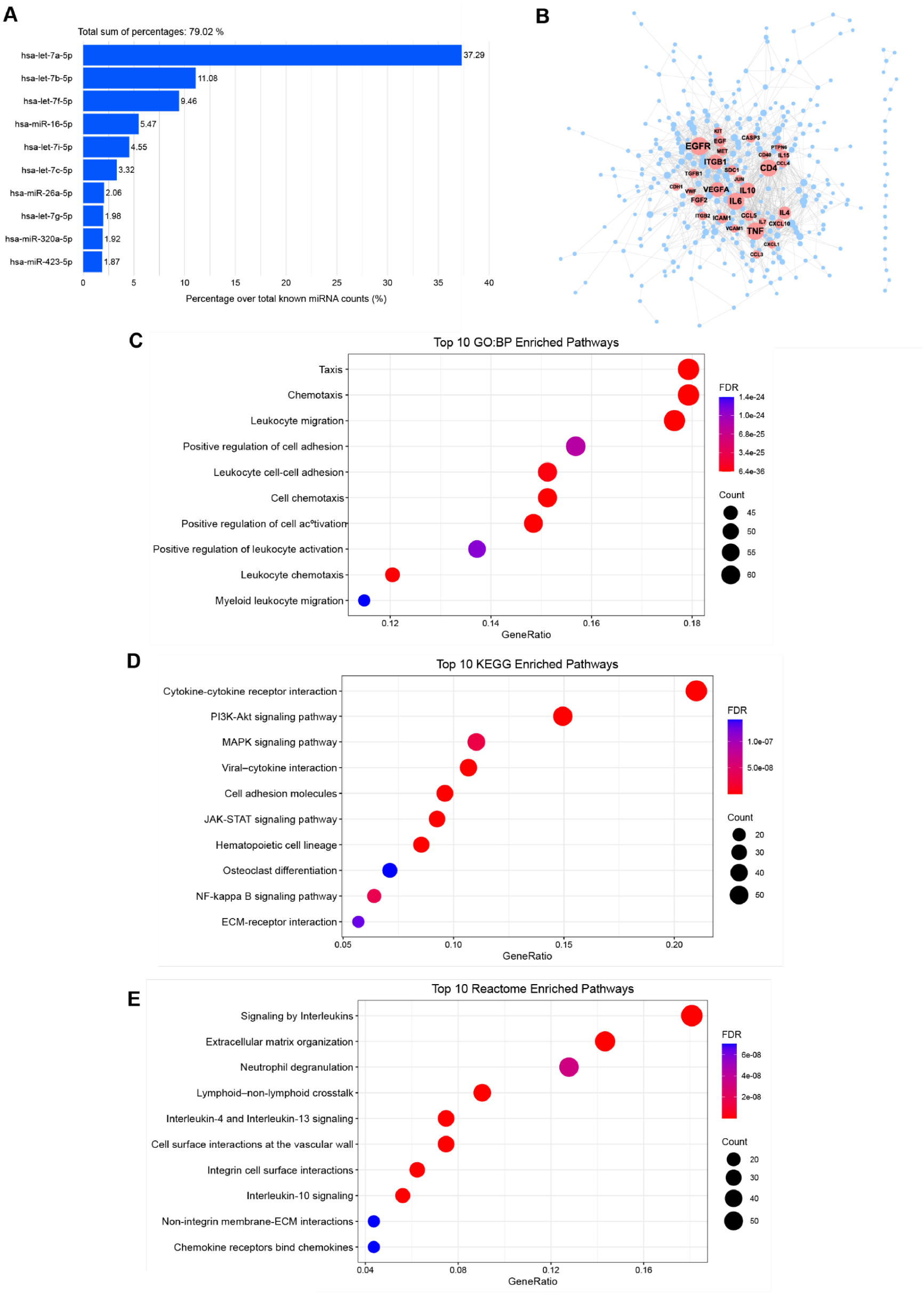
PRS CK STORM is a complex biologic rich in immunomodulating miRNAs and proteins. (**A**)Top 10 most abundant exosomal microRNAs in PRS CK STORM. Exosomal miRNA profiling was performed on five biological replicates of PRS CK STORM. Total RNA was extracted from 2 mL of each secretome. The relative abundance of each miRNA species was calculated independently per sample, and the mean abundance across the five replicates was used to rank miRNAs by overall representation. (**B**)High-confidence protein–protein interaction (PPI) network of the PRS CK STORM proteome. PPI network constructed from 362 proteins detected above the assay-specific limit of detection in at least three of five biological replicates and exhibiting at least one high-confidence interaction (combined STRING score ≥ 0.7). The network was visualized using an organic layout in Cytoscape (v3.10.2). Node and label sizes were scaled by degree centrality; only nodes with degree ≥ 20 are labeled. The resulting network reveals a densely interconnected secretome architecture consistent with coordinated immunoregulatory and tissue repair functions. (**C, D, E**) Functional enrichment analysis of the PRS CK STORM protein interaction network. Overrepresented biological functions were identified from the high-confidence interaction network (n=362 proteins) using three curated resources: Gene Ontology Biological Processes (GO:BP), KEGG, and Reactome. (**C**) Top 10 most significantly enriched GO:BP terms highlight roles in immune activation, leukocyte migration, and cell adhesion. (**D**) Top 10 most significantly enriched KEGG pathways comprise canonical inflammatory and proliferative signaling (e.g., JAK–STAT, NF-κB, PI3K–Akt, MAPK), ECM remodeling, and host–pathogen interactions. (**E**) Top 10 most significantly enriched Reactome results identify anti-inflammatory cytokine signaling (IL-10, IL-4, IL-13), lymphoid– non-lymphoid cell communication, and ECM reorganization via integrin and non-integrin interfaces. Only pathways with FDR < 0.05 are shown.

### Targeted Proteomics Uncovers a Core Immunoregulatory Network in PRS CK STORM

To further characterize the composition of PRS CK STORM, we carried out a targeted proteomic analysis using the Olink Explore 3072 assay. A total of 498 proteins were detected above the assay-specific limit of detection (LOD) in at least three of the five biological replicates of PRS CK STORM (Table S4). To explore potential functional associations, this protein set was used to construct a protein–protein interaction (PPI) network using the STRING database. To retain biologically meaningful connectivity, only proteins with at least one high-confidence interaction (interaction score ≥ 0.7) were included. The resulting network comprised 362 proteins (Table S5) and revealed a densely interconnected architecture, consistent with a coordinated secretome supporting immunoregulatory and tissue repair functions (Figure 3B). Topological analysis (based on degree and betweenness centrality) identified several high-scoring hub proteins, including epidermal growth factor receptor (EGFR), vascular endothelial growth factor A (VEGFA), fibroblast growth factor 2 (FGF2), and transforming growth factor beta 1 (TGFB1). These molecules are central mediators of epithelial regeneration, extracellular matrix (ECM) remodeling, and angiogenesis (31–34). Such factors may have contributed to the angiogenesis observed in the lung tissue of mice treated with PRS CK STORM in our *in vivo* Sars-CoV-2 model. Anti-inflammatory activity was reflected by the presence of canonical immune modulators IL-10 and IL-4, both of which facilitate resolution of inflammation (35,36), with the former specifically involved in inhibiting IFN-γ (37). CD4, a core marker of helper T cell identity and function (38), also appeared as a central node, underscoring the complex immunomodulatory composition of PRS CK STORM.

Additional central nodes included adhesion- and ECM-associated proteins such as intercellular adhesion molecule 1 (ICAM1), vascular cell adhesion molecule 1 (VCAM1), and integrins beta-1 and beta-2 (ITGB1, ITGB2), supporting functional roles in leukocyte trafficking and tissue structural integrity (35,39). Several chemokines, including CCL3, CCL4, CCL5, and C-X-C motif chemokine ligand 10 (CXCL10), were also prominent, reflecting a capacity to coordinate immune cell recruitment (40). Furthermore, classical pro-inflammatory mediators such as TNF-α, IL6, and IL-15 emerged as hubs, reflecting their involvement in immune signaling cascades (35,36).

The systems-level functions of the PRS CK STORM proteome were elucidated through functional enrichment analysis of the entire interaction network using GO:BP, KEGG, and Reactome databases (Figures 3C, D, E). GO:BP terms revealed significant enrichment in processes related to immune cell activation, leukocyte trafficking, and adhesion. KEGG pathway analysis further highlighted canonical inflammatory and proliferative signaling cascades (JAK–STAT, NF-κB, PI3K–Akt, MAPK signaling), alongside ECM remodeling and host–pathogen interaction pathways. Reactome analysis expanded on these findings, identifying key pathways involved in immunoregulation and structural remodeling, including anti-inflammatory cytokine signaling (IL-10, IL-4, IL-13), lymphoid– non-lymphoid cell communication, and ECM organization via integrin and non-integrin interfaces. Full enrichment results are provided in Supplemental Tables S6A, B, C. Overall, the proteomic profile of PRS CK STORM reveals a concerted enrichment in pathways governing immune regulation, extracellular matrix remodeling, and inflammatory resolution.

## Discussion

ARDS is a serious health issue for which effective pharmacological therapies are limited (2). In this study, we demonstrate that PRS CK STORM represents a compelling therapeutic strategy for treating cytokine storm–driven ARDS, via a multimodal mechanism of action that integrates broad immunomodulation with tissue repair.

### Therapeutic Efficacy Compared to Corticosteroids

The standard method of care for COVID-19 patients requiring respiratory support is treatment with the corticoid dexamethasone (8). While effective in reducing mortality, dexamethasone can cause a variety of severe adverse reactions including hyperglycemia, gastrointestinal hemorrhage, and psychosis (9). Perhaps even more concerning, in the context of respiratory viral infections, treatment with corticoids can cause immunosuppression, leading to slower viral clearance and risk of secondary infections (8–10). In our lethal SARS-CoV-2–induced ARDS model, PRS CK STORM was at least as effective as dexamethasone in ameliorating histopathological lung damage. The relatively high number of pulmonary blood capillaries in PRS CK STORM-treated mice, surpassing that of corticoid-treated mice, is indicative of angiogenesis-mediated tissue repair (41). Previous work by our group demonstrated that PRS CK STORM has no serious adverse effects in an *in vivo* mouse model (18). Altogether, these data position PRS CK STORM as an equally potent alternative to corticoids that is potentially safer and has complementary mechanisms for treatment of ARDS.

### Mechanistic Insights: Macrophage Reprogramming, COX2 Inhibition, Immunomodulation Factors

Our *in vitro* data clearly demonstrates the immunomodulatory effects of PRS CK STORM are at least in part due to its repolarization of pro-inflammatory macrophages toward a reparative phenotype. In LPS-stimulated THP-1 derived macrophages, transcriptomic profiling revealed PRS CK STORM selectively downregulates central nodes of the TLR signaling cascade, including *MyD88, TRAF6, IKK2, NFKB1*, and *COX2*. This builds on our previous work that showed substantial reduction of levels of prototypical pro-inflammatory cytokines like TNF-α and IL-1β (10,11). Interestingly, we found that PRS CK STORM downregulates *COX2* not only on the level of gene expression but also by directly inhibiting its enzymatic activity, suggesting this secretome is a multivalent biologic. Thus, enzymatic assays revealed potent, dose-dependent inhibition of COX2 activity by PRS CK STORM. This inhibitory effect was consistent across five independent GMP-grade production lots, confirming batch-to-batch reproducibility. Given that COX2 plays a central role in the synthesis of pro-inflammatory prostaglandins (42), its inhibition represents a key mechanism of PRS CK STORM’s anti-inflammatory action.

Lending weight to the idea that PRS CK STORM is a multivalent biologic, we have found that this secretome has a complex and varied composition. While causality between individual miRNAs and observed effects cannot be established from abundance alone, the exosomal miRNAs found in this secretome are consistent with an anti-inflammatory profile, with an overrepresentation of the let-7 family and other miRNAs involved in inflammation resolution and tissue repair. Proteomic analyses showed elevated levels of endogenous regulators known to attenuate inflammation such as IL-10 and IL-4, as well as regenerative mediators such as EGFR, VEGFA, FGF2, and TGFB1. It is important to note that such elevated levels do not necessarily reflect high protein abundance in PRS CK STORM, as the Olink Normalized Protein eXpression (NPX) metric represents relative expression and cannot be used to infer absolute concentration levels.

The detection of pro-inflammatory or chemotactic proteins within the PRS CK STORM secretome should not be misconstrued as indicative of a detrimental or undesired profile. Rather, it reflects the physiological complexity of the immunological crosstalk between M2 macrophages and MSCs. The detection of certain inflammatory mediators likely mirrors their roles as transient yet essential regulators of inflammation resolution and tissue repair. For example, TNF-related apoptosis-inducing ligand (TRAIL) has been shown to promote neutrophil apoptosis and accelerate the resolution of inflammation in both acute lung injury and peritonitis models (43). Similarly, IL-6, through trans-signaling mechanisms mediated by soluble IL-6 receptor (sIL6R) released from apoptotic neutrophils, governs the recruitment of monocytes essential for the clearance of dying cells and transition to tissue repair (44). These findings emphasize that the presence of such factors in PRS CK STORM reflects a coordinated, resolution-oriented immune signature, rather than persistent inflammatory activation. Altogether, the molecular composition of PRS CK STORM is coherent with its *in vivo* efficacy in our ARDS murine model and complements mechanistic insights from the *in vitro* transcriptomic assays, reinforcing its potential to modulate key checkpoints in immune homeostasis.

### Resolution Pharmacology and Translational Outlook

PRS CK STORM exemplifies the emerging paradigm of resolution pharmacology: multi-target interventions aimed at restoring homeostasis of the immune system rather than overly suppressing it (12,45). Our data strongly suggest that, unlike conventional therapies that interrupt signaling downstream or neutralize single cytokines, PRS CK STORM exerts a multi-target immunomodulation with the added benefit of supporting tissue repair, offering a novel therapeutic strategy for cytokine storm–driven ARDS (Figure 4). Beyond its demonstrated efficacy in viral ARDS, PRS CK STORM emerges as a platform immunotherapy with broad potential across macrophage-driven hyperinflammatory syndromes. Its dual action—coordinated immunomodulation and regenerative signaling—positions it to address cytokine release syndrome (CRS) in CAR-T therapy, secondary hemophagocytic lymphohistiocytosis (HLH), macrophage activation syndrome (MAS), and autoimmune flare states like systemic lupus erythematosus (46– 50). In these settings, PRS CK STORM could suppress upstream inflammatory drivers such as monocyte/macrophage-derived IL-6, IFN-γ, and TNF-α without compromising immune competence, while promoting tissue repair through factors like VEGFA, TGFB1, and EGFR.

**Figure 4.**
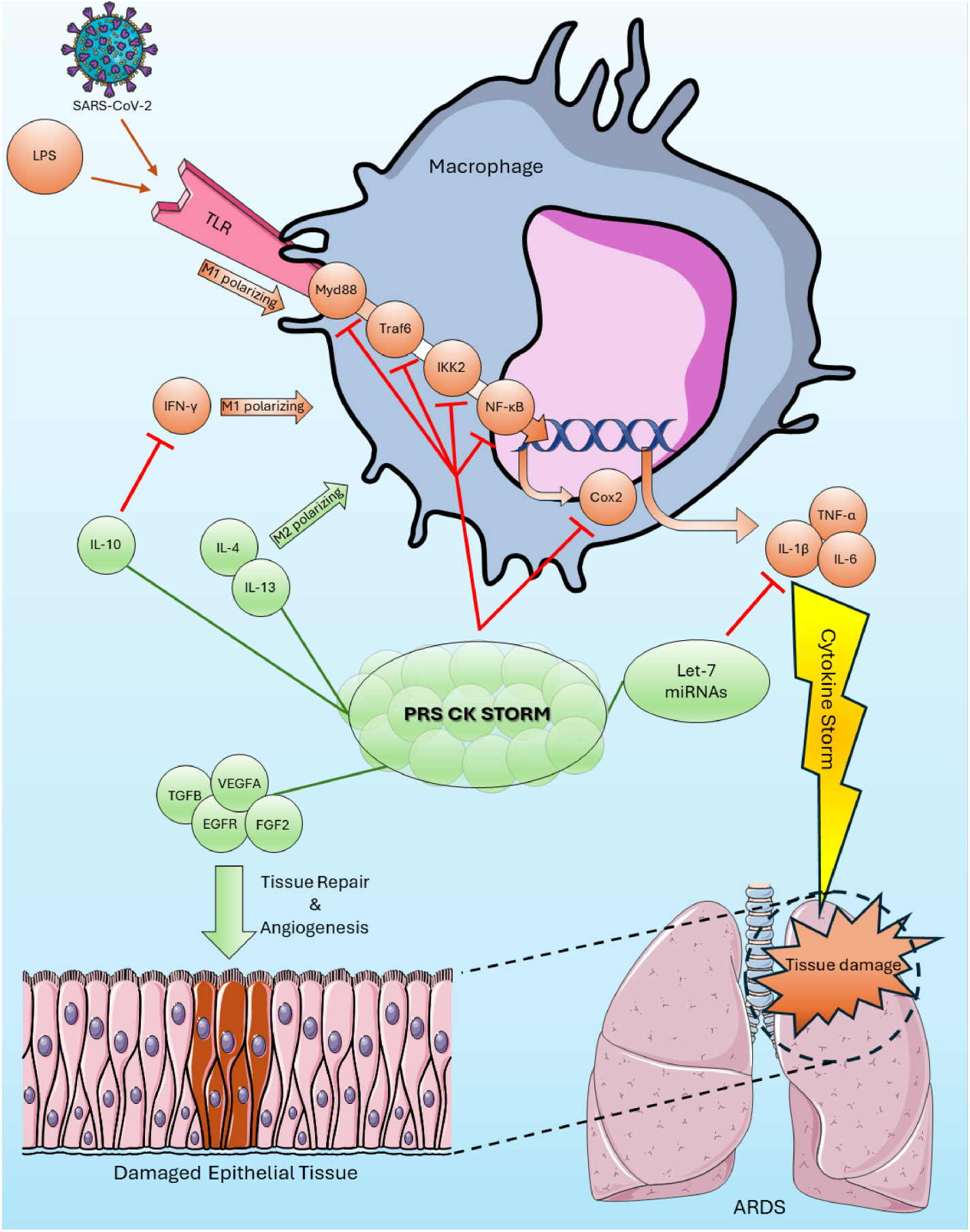
Working model for the mechanism of action of PRS CK STORM in the context of ARDS. TLRs in macrophages are activated via a PAMP such as SARS-CoV-2 or LPS, triggering a signaling cascade that includes Myd88, Traf6, IKK2, and NF-ĸB, ultimately inducing the expression of pro-inflammatory enzyme COX2 and pro-inflammatory cytokines such as TNF-α, IL-1β, and IL-6. This, combined with the pro-inflammatory activity of other cytokines such as IFN-γ, leads to the M1 polarization of macrophages. When macrophages become locked in this state, they produce increasing numbers of pro-inflammatory cytokines, leading to cytokine storm, ARDS, and damage to lung tissue. PRS CK STORM is a multivalent biologic that counteracts this detrimental cascade via a variety of avenues. M2-polarizing cytokines found in this secretome such as IL-4 and IL-13 reduce the pro-inflammatory state of macrophages, while IL-10 inhibits the effects of IFN-γ. PRS CK STORM directly inhibits the activity of the COX2 enzyme and downregulates the expression of several key components of the pro-inflammatory TLR pathway (Myd88, Traf6, IKK2, NF-ĸB). Further downstream, the let-7 family of miRNAs downregulates pro-inflammatory cytokines TNF-α, IL-1β, and IL-6. In addition to curtailing the immune response, PRS CK STORM promotes angiogenesis and healing in damaged tissue via regenerative factors such as TGFB, VEGFA, EGFR, and FGF2.

The consistent efficacy of PRS CK STORM across five independent GMP batches underscores the industrial robustness of the co-culture platform, and its cell-free nature indicates a broad safety profile. Although this early preclinical evidence supports the therapeutic potential of an MSC-derived secretome in ARDS and cytokine storm syndromes, robust clinical validation is essential to confirm its translational feasibility. Our study is limited by its preclinical design and the use of a single viral strain. Future studies should assess pharmacokinetics, dose-response relationships, and examine efficacy in large animals. Additionally, future trials should explore its application across diverse immunopathologies.

## Conclusions

These data position PRS CK STORM as a next-generation, cell-free immunoregenerative biologic capable of attenuating hyperinflammation and promoting lung repair in viral ARDS in a manner reminiscent of physiological inflammation resolution. Given its multimodal mechanism, batch reproducibility, and off-the-shelf applicability, PRS CK STORM may serve as a platform therapy for cytokine storms across diverse etiologies, viral or otherwise, enabling widespread immune recalibration where conventional single-target interventions fall short.

## Supporting information

Supplementary methods, tables, references

Supplementary excel tables for proteomics

## Abbreviations

APC: adenomatosis polyposis coli
ARDS: acute respiratory distress syndrome
BIPF: bacteria-induced pulmonary fibrosis
CCL: chemokine ligand
CCL2: C-C motif chemokine ligand 2
COX2: cyclooxygenase-2
COVID-19: coronavirus disease 19
CRS: cytokine release syndrome
Ct: cycle threshold
CXCL10: C-X-C motif chemokine ligand 10
DAMPs: damage-associated molecular patterns
ECM: extracellular matrix
EGFR: epidermal growth factor receptor
FGF2: fibroblast growth factor 2
GM-CSF: granulocyte-macrophage colony-stimulating factor
GMP: good manufacturing practices
gp130: glycoprotein 130
HGF: hepatocyte growth factor
HLH: hemophagocytic lymphohistiocytosis
ICAM1: intercellular adhesion molecule 1
ICU: intensive care unit
IFN-γ: interferon gamma
IGF: insulin-like growth factor
IKK2: IκB kinase 2
IL: interleukin
IL-1β: interleukin-1 beta
IL-1Ra: interleukin-1 receptor antagonist
IQR: interquartile range
ITGB1: integrin beta-1
ITGB2: integrin beta-2
JAK–STAT: janus kinase–signal transducer and activator of transcription
LOD: limit of detection
LPS: lipopolysaccharide
MAPK: mitogen-activated protein kinase
MAS: macrophage activation syndrome
miRNA: microRNAs
MSCs: mesenchymal stromal cells
MYD88: myeloid differentiation primary response 88
NF-κB: nuclear factor kappa-light-chain-enhancer of activated B cells
NFKB1: NF-κB subunit 1
NPX: Normalized Protein eXpression
PAMPs: pathogen-associated molecular patterns
PFU: plaque-forming units
PI3K: phosphoinositide 3-kinase
PPI: protein–protein interaction
PRRs: pattern recognition receptors
sIL6R: soluble IL-6 receptor
SOD2: superoxide dismutase 2
STAT3: signal transducer and activator of transcription 3
TGFB1: transforming growth factor beta 1
THP-1: Tohoku hospital pediatrics-1
TLR: toll-like receptor
TNF-α: tumor necrosis factor-alpha
TRAF6: TNF receptor associated factor 6
TRAIL: TNF-related apoptosis-inducing ligand
VCAM1: vascular cell adhesion molecule 1
VEGFA: vascular endothelial growth factor A

## Availability of data and materials

Both raw and processed microRNA sequencing data have been deposited in the Gene Expression Omnibus under accession number GSE301236. https://www.ncbi.nlm.nih.gov/geo/

## Authors’ contributions

Author Contributions: This study was conceived, designed, and supervised by JC.DG, F.S, and JP.L. In vivo experiments were performed by JA.Q, J.S, P.F, K.C, and P.D, with analysis and interpretation carried out by C.P.L, AR.GR, and C.CT. In vitro experiments were performed by N.MC, B.G, A.DG and analyzed by N.MC, J.GM, and F.OG. Bioinformatics analyses and interpretation were caried out by D.L. The manuscript was written by C.P.L, D.L, and JP.L, and revised and approved by all authors.

## Acknowledgements

The authors thank Luis F. de Villa Alcázar (Hospital Ruber Internacional) for critical reading of the manuscript and Melchor Alvarez de Mon (Hospital Universitario Príncipe de Asturias) for his expert technical assistance. We also thank the Hospital of Fuenlabrada for the use of their facilities.

## Supplementary Information

Supplementary Material 1.docx

Contains supplementary methods, tables, and references.

Supplementary Material 2.xlsx

Contains large supplementary tables related to the proteomics analysis.

