## Supplementary methods, tables, references for "A Novel Secretome Rewrites the Immune Response in Viral Acute Respiratory Distress Syndrome"

#### *In vivo* SARS-CoV-2 ARDS Model and Treatment: Extended

A total of 40 male B6.Cg-Tg(K18-ACE2)2Prlmn/J or K18-hACE2 transgenic mice (Stock No: 034860) from Jackson Laboratories (https://www.jax.org/strain/034860) were used. These mice express the human ACE2 protein that acts as a receptor for SARS-CoV-1-2002 and SARS-CoV-2-2019, under the human cytokeratin 18 (K18) promoter, which directs the expression of the transgene to the respiratory tract.

The virus used in the experiment was SARS-CoV-2 strain Slovakia/SK-BMC-BA43/2022- Omicron VOC BE.1.1 (aka BA.5.3.1.1.1.1.) (006V-04753). The dose used ($10^5$ pfu in intranasal instillation), is a 100 lethal dose (LD100), and it has been demonstrated in scientific literature to generate a very aggressive cytokine storm model and a strong cerebral oedema caused by the large viral replication that occurs in the brain, leading to near uniform death by day 8 (1).

Animals were allocated into 4 groups as follows:

- **Negative control group:** 3 genetically modified h-ACE2 mice without infection or treatment (mice 1.2, 1.3, 1.4).
- **Positive control group:** 12 genetically modified h-ACE2 mice with intranasal infection of $10^5$ pfu of SARS-CoV-2 supplied by INIA-CSIC of Valdeolmos. This group was not treated at any time with any type of drug (mice 2.1, 2.2, 2.3, 2.4, 3.1, 3.2, 3.3, 3.4, 4.1, 4.2, 4.3, 4.4).
- **Corticoid group:** 12 genetically modified h-ACE2 mice with intranasal infection of $10^5$ pfu of SARS-CoV-2 supplied by INIA-CSIC of Valdeolmos. This group was treated intravenously with 40 µL of dexamethasone dosed at 5mg/kg, for 5 consecutive days (Days 1, 2, 3, 4 and 5 of the experiment) (mice 5.1, 5.2, 5.3, 5.4, 6.1, 6.2, 6.3, 6.4, 7.1, 7.2, 7.3, 7.4).
- **PRS CK STORM group:** 12 genetically modified h-ACE2 mice with intranasal infection of $10^5$ pfu of SARS-CoV-2 supplied by INIA-CSIC of Valdeolmos. This group was treated intravenously with 40 µL of PRS CK STORM (containing 11,897 pg of TIMP-1), for 5 consecutive days (Days 1, 2, 3, 4 and 5 of the experiment) (mice 8.1, 8.2, 8.3, 8.4, 9.1, 9.2, 9.3, 9.4, 10.1, 10.2, 10.3, 10.4). This dose was based on (2).

The trial was performed double-blind, with a total duration of 8 days. Day 0 was collection of initial baseline data and subsequent inoculation of the virus in the mice of the infected groups. On day 1 (24h post infection), animals in treatment groups received an IV injection of the drug, dexamethasone or PRS CK STORM. 1 minute later, blood was extracted to perform the analysis of murine IFN-γ. On days 2, 3, 4, 5 the same treatments as on day 1 were repeated, with no interventions on days 6, 7. On day 8, blood extractions were repeated. All animals were sacrificed by day 8. Throughout the study, we sought to respect as much as possible the 3R principle of animal experimentation. When an animal's weight fell below 75%, or the animal exhibited strong signs of discomfort, it was sacrificed prematurely and the final exsanguination and removal of the lungs for histopathology and viral load studies were carried out. To verify that the viral load had been effective in generating the model ($10^5$ pfu by intranasal instillation), the viral load was analyzed by studying the cycle threshold (Ct) (3). Sections of lung tissue were subjected to hematoxylin and eosin staining for microscopic inspection. The serum concentration of murine IFN-γ was quantified by multiplex ELISA.

#### Exosomal RNA Extraction, MicroRNA Library Preparation and Sequencing

Total exosome-derived RNA was extracted from 2 mL of PRS CK STORM using the exoRNeasy Maxi Kit (QIAGEN), following the manufacturer's instructions, and eluted in 14 µL. Small RNA libraries were generated from 5 µL of total RNA using the QIAseq miRNA Library Kit (QIAGEN). Unique molecular identifiers (UMIs) were incorporated during reverse transcription after adapter ligation. Complementary DNA was amplified by PCR (22 cycles), during which sample indices were added. Amplified products were purified and assessed for quality and fragment size using capillary electrophoresis (TapeStation D1000). Libraries were pooled at equimolar concentrations based on insert size and concentration, quantified by qPCR, and sequenced on a NextSeq 2000 instrument using a 1 × 75 bp and 2 × 10 bp configuration, according to the manufacturer's protocol. Exosomal microRNA profiling was performed on five biological replicates of PRS CK STORM. Raw reads were demultiplexed and FASTQ files generated for each sample.

### MicroRNA Data Processing and Analysis

Sequencing quality was assessed using FastQC (v0.11.9) (4). Adapter trimming, quality filtering, and read-length selection were performed with Trim Galore! (v0.6.10) (5) using parameters optimized for human microRNA size. UMIs were extracted and PCR duplicates removed using the extract and dedup modules of UMI-tools (v1.1.5) (6). Deduplicated reads were aligned to the human reference genome (GRCh38) using miRDeep2 (v2.0.1.2) (7). Quantification of known microRNA species was performed using the miRDeep2 quantifier.pl module, with mature and hairpin sequences for *Homo sapiens* and *Mus musculus* retrieved from miRBase (release 22.1) used as reference input (8).

The relative abundance of each microRNA species was calculated independently per sample, and the mean abundance across the five replicates was used to rank microRNAs by overall representation. The top 10 most abundant exosomal microRNAs—collectively accounting for nearly 80% of the total profile—were selected for functional annotation, based on the rationale that dominant species are most likely to mediate biologically meaningful effects.

Both raw and processed microRNA sequencing data have been deposited in the Gene Expression Omnibus (9,10) under accession number GSE301236 .

### Olink Proteomic Assay and Data Preprocessing

Proteomic profiling of PRS CK STORM was performed on five biological replicates by Health in Code (Spain), a certified provider of the Olink Explore platform. Protein abundance was quantified using the Olink Explore 3072 assay, which combines antibody-based detection with next-generation sequencing (NGS) readout on the NovaSeq 6000 system (Illumina). Sample preparation and assay execution were conducted according to the manufacturer's protocol (Olink Explore, Sweden). Briefly, serum samples were diluted to optimal conditions and incubated in 384-well plates with dual oligonucleotide-labeled antibody probes. Upon binding to their target proteins, proximity extension and PCR amplification steps generated protein-specific DNA barcodes, which were pooled into indexed libraries and sequenced.

The present analysis focused on proteins measured within the Inflammatory I and Cardiometabolic I panels. Raw data were processed using Olink's internal quality control pipeline, and proteins failing batch release criteria were excluded. Expression levels were reported as Normalized Protein eXpression (NPX) values—arbitrary $log_2$-scaled units derived from quantification cycle (Cq) values. NPX normalization accounts for intra- and inter-assay variability, allowing relative comparisons within a given protein across samples. NPX values cannot be used to compare absolute abundance between different proteins. A one-unit change in NPX corresponds to a two-fold change in relative expression. Only proteins detected above the assay-specific LOD in at least three biological replicates were retained for downstream analysis.

### Protein–Protein Interaction and Network Topology Analysis

To explore functional relationships within the PRS CK STORM proteome, a protein–protein interaction (PPI) network was constructed using the STRING database (version 11.5) (11), applying a high-confidence interaction threshold (combined score ≥ 0.7). Only proteins with at least one reported interaction (edge) were retained in the network. All analyses were performed in R (v4.4.2) (12). Network analysis and data manipulation were conducted using STRINGdb (v2.18.0) (11), igraph (v2.1.4) (13), and dplyr (v1.1.4) (14).

The interaction network was visualized in Cytoscape (v3.10.2) (15) using an organic force-directed layout. Topological properties, including node degree and betweenness centrality, were computed with NetworkAnalyzer plugin (16) to identify highly connected hub proteins and potential regulatory intermediates. In the final visualization, node and label sizes were scaled according to degree centrality, with labels displayed only for proteins exhibiting a degree ≥ 20.

### Functional Enrichment Analysis

Functional enrichment analysis was conducted to identify biological pathways and processes overrepresented in the PRS CK STORM proteome. Gene Ontology Biological Processes (GO:BP) (17), Kyoto Encyclopedia of Genes and Genomes (KEGG) pathways (18), and Reactome pathways (19) were analyzed using the clusterProfiler (v4.14.4) (20) and ReactomePA (v1.50.0) (21) R packages. Protein identifiers were mapped to Entrez Gene IDs using the org.Hs.eg.db annotation package (v3.20.0) (22). Overrepresentation testing was performed using one-sided Fisher's exact tests, with multiple comparisons corrected via the Benjamini–Hochberg procedure. Pathways with a false discovery rate

(FDR) < 0.05 were considered statistically significant. Enrichment results were visualized with ggplot2 (v3.5.1) (23), and composite figure panels were assembled using cowplot (v1.1.3) (24).

### Supplementary Tables

**Table S1.** List of primers used in qPCRs to measure relative gene expression.

| **Primer** | **Sequence 5' to 3'** |
|---|---|
| *TRAF6* Forward | TTGTCCACACAATGCAAGGAG |
| *TRAF6* Reverse | TGGCGTCCATGACCTCTTC |
| *IKK2* Forward | ACAGCGAGCAAACCGAGTTTGG |
| *IKK2* Reverse | CCTCTGTAAGTCCACAATGTCGG |
| *MYD88* Forward | GAGGCTGAGAAGCCTTTACAGG |
| *MYD88* Reverse | GCAGATGAAGGCATCGAAACGC |
| *NFKB1* Forward | CTGGTGCATTCTGACCTTGC |
| *NFKB1* Reverse | GGTCCATCTCCTTGGTCTGC |
| *NLRP3* Forward | GGACTGAAGCACCTGTTGTGCA |
| *NLRP3* Reverse | TCCTGAGTCTCCCAAGGCATTC |
| *COX2* Forward | CGGTGAAACTCTGGCTAGACAG |
| *COX2* Reverse | GCAAACCGTAGATGCTCAGGGA |

**Table S2.** Protocol for assigning lung histopathological scores for each measured parameter.

| | Degree of Alteration | | | | |
|---|---|---|---|---|---|
| **Parameter** | 0 | 1 | 2 | 3 | 4 |
| Proliferation of ATII cells | No | Scattered | Few | Moderate | Moderate-intense |
| Interstitial Inflammation | No | Scattered cells | 1 layer | 2 layers | >2 layers |
| Peribronquiolar Cuffing (Mononuclear cells) | No | Scattered cells | 1 layer | 2 layers | >2 layers |
| Perivascular edema | No | Mild | Moderate | Moderate to intense | Intense |
| Alveolar edema + cells | No | Mild edema | Mild edema 1-2 cells | Moderate edema 4-6 cells | Moderate edema >6 cells |

**Table S3.** Top 10 most abundant exosomal microRNAs in PRS CK STORM and their reported functional associations in respiratory and inflammatory disease contexts.

| **MicroRNAs** | **Mean % of Total Abundance** | **Reported Functional Associations*** |
|---|---|---|
| hsa-let-7a-5p | 37.29% | Negatively regulates pro-inflammatory cytokines IL-6, TNF-α, IL-1β and Ras–MAPK signaling in airway inflammation (25); promotes anti-fibrotic effects, reduced macrophage infiltration, IL-10 secretion in lung injury models (26) |

| | | |
|---|---|---|
| hsa-let-7b-5p | 11.08% | Reduces TLR4/NF-κB signaling and lung injury in sepsis models (27); linked to interferon regulation and antiviral defense during influenza infection (28) |
| hsa-let-7f-5p | 9.46% | Negatively regulates PI3K/Akt/COX2 signaling in BIPF animal models (29); suppresses NLRP3 inflammasome activation in BMSCs derived from SLE patients (30) |
| hsa-miR-16-5p | 5.47% | Suppresses TXNIP and inhibits NLRP3 inflammasome activation in macrophages derived from CHD patients (31); restores Th17/Treg balance and modulates cytokine expression via LATS1 targeting in T cells from SLE patients (32) |
| hsa-let-7i-5p | 4.55% | Inhibits activation of fibroblasts through TGFBR1/Smad3 signaling modulation, alleviating pulmonary fibrosis (33) |
| hsa-let-7c-5p | 3.32% | Inhibits M2 macrophage polarization and MMP-9/12 release via the IL-6/STAT3 pathway in airway epithelial cells, decreasing COPD emphysema (34) |
| hsa-miR-26a-5p | 2.06% | Decreases pro-inflammatory responses in ARDS murine models by Wnt/β-catenin pathway activation (35) and CHUK/NF-κB pathway inhibition (36) |
| hsa-let-7g-5p | 1.98% | Inhibits Treg-to-Th17 cell transdifferentiation by suppressing STAT3 expression in collagen-induced arthritis mice models (37) |
| hsa-miR-320a-5p | 1.92% | Modulates collagen production via TGFBR2 and IGF1R; dysregulated in fibrosis in SSc-ILD (38) |
| hsa-miR-423-5p | 1.87% | Attenuates RV-induced inflammatory injury and NLRP3 inflammasome activation in a murine asthma model by |

| | | |
|---|---|---|
| | | targeting PINK1 (39); proposed as a potential diagnostic biomarker in COPD (40) |

* Functional associations are drawn from published studies in respiratory and inflammatory disease models. Given the pleiotropic and context-specific nature of microRNA activity, functions may vary across biological contexts.

*Definition of abbreviations*: IL = interleukin; TNF-α = tumor necrosis factor alpha; MAPK = mitogen-activated protein kinase; PI3K = phosphoinositide 3-kinase; Akt = protein kinase B; COX2 = cyclooxygenase-2; BIPF = bacteria-induced pulmonary fibrosis; NLRP3 = NOD-like receptor family, pyrin domain containing 3; BMSCs = bone marrow–derived mesenchymal stem cells; SLE = systemic lupus erythematosus; TLR4 = Toll-like receptor 4; NF-κB = nuclear factor kappa B; TXNIP = thioredoxin-interacting protein; CHD = coronary heart disease; Th17 = T helper 17 cells; Treg = regulatory T cells; LATS1 = large tumor suppressor kinase 1; TGFBR = transforming growth factor beta receptor; Smad3 = SMAD family member 3; MMP = matrix metalloproteinase; STAT3 = signal transducer and activator of transcription 3; COPD = chronic obstructive pulmonary disease; ARDS = acute respiratory distress syndrome; CHUK = conserved helix-loop-helix ubiquitous kinase; IGF1R = insulin-like growth factor 1 receptor; SSc-ILD = systemic sclerosis–associated interstitial lung disease; RV = rhinovirus; PINK1 = phosphatase and tensin homolog-induced putative kinase 1.

**Table S4.** Proteins Detected Above the Assay-Specific Limit of Detection in PRS CK STORM Across Biological Replicates.

[See external excel spreadsheet: Table S4]

**Table S5.** Proteins Retained in the PRS CK STORM Interaction Network and their Topological Properties.

[See external excel spreadsheet: Table S5]

**Table S6.**

**(A)** Gene Ontology Biological Processes (GO:BP) Enrichment of the PRS CK STORM Interaction Network.

[See external excel spreadsheet: Table S6A]

**(B)** KEGG Pathway Enrichment of the PRS CK STORM Interaction Network.

[See external excel spreadsheet: Table S6B]

**(C)** Reactome Pathway Enrichment of the PRS CK STORM Interaction Network.

[See external excel spreadsheet: Table S6C]
